# Decomposing the Apoptosis Pathway Into Biologically Interpretable Principal Components

**DOI:** 10.1101/237883

**Authors:** Min Wang, Steven M. Kornblau, Kevin R. Coombes

## Abstract

Principal component analysis (PCA) is one of the most common techniques in the analysis of biological data sets, but applying PCA raises two challenges. First, one must determine the number of significant principal components (PCs). Second, because each PC is a linear combination of genes, it rarely has a biological interpretation. Existing methods to determine the number of PCs are either subjective or computationally extensive. We review several methods and describe a new R package, **PCDimension**, that implements additional methods, the most important being an algorithm that extends and automates a graphical Bayesian method. Using simulations, we compared the methods. Our newly automated procedure performs best when considering both accuracy and speed. We applied the method to a proteomics data set from acute myeloid leukemia patients. Proteins in the apoptosis pathway could be explained using six PCs. By clustering the proteins in PC space, we were able to replace the PCs by six “biological components”, three of which could be immediately interpreted from the current literature. We expect this approach combining PCA with clustering to be widely applicable.

## Introduction

Since the earliest days of gene expression microarrays, two-way clustered heatmaps have been a standard feature of papers studying genome-wide biological data sets.^1,2^ Such heatmaps remain ubiquitous, in spite of numerous difficulties in interpretation, reproducibility, and in assigning statistical significance. Good clustering of the genes in such data sets is critical for understanding the biology. Because many biologists are more interested in signaling pathways than in individual genes, they want to find a source of consistent, robust, and interpretable blocks of genes that drive distinct functional characteristics of the pathways. These blocks of genes form clusters that are relevant to comprehensive understanding of critical biological processes. For example, apoptosis is an important biological process, which is characterized by distinct morphological states and energy-dependent biochemical mechanisms.^3^ Understanding how proteins cluster in the apoptotic pathway will help elucidate its underlying molecular mechanisms. Now, clustering can be thought of as a form of dimension reduction, and a natural question is the “true dimension” of the data. Various techniques have been developed to determine the dimensionality, the most common being principal component analysis (PCA). For our purposes, an important problem in PCA is to determine the number of statistically significant components.

Numerous methods have already been developed to estimate the number of significant components. There are three main types of approaches: (1) *ad hoc* subjective and graphical rules, (2) methods based on distributional assumptions, and (3) computationally extensive procedures relying on Monte Carlo, permutation, cross-validation, bootstrap, or jackknife.^4^ The screeplot method, which consists of plotting a curve of the eigenvalues of the sample covariance matrix versus their rank and looking for an “elbow” in the curve, is the most famous graphical approach.^5^ However, this method relies on the user’s subjective experience to find any possible “elbow”. Even so, other methods are not always superior to the simple screeplot. Legendre and Legendre used the “broken stick” distribution to compare the extra information in a model to one with fewer parameters.^6^ Ferre carried out an empirical study of many methods to select the number of PCs, using data simulated from known parameters.^7^ He concluded that there is no “ideal” solution to the problem of dimensionality in PCA. He also concluded that Bartlett’s tests^8^ are an improvement because they are less subjective, but may have a tendency to overestimate the true number of components. More recently, Peres-Neto and colleagues conducted an extensive simulation study to evaluate a wider variety of methods.^9^ They concluded that several methods, especially those based on randomization and permutation proposed by ter Braak,^10,11^ outperform the others and should be applied to study general data sets.

In 2008, Auer and Gervini addressed the problem of selecting principal components in the context of Bayesian model selection.^12^ While their method has strong theoretical foundations and appears to work well in practice, it still depends on the subjective evaluation of a graphical display of how the maximum posterior estimate of the number of components depends on a parameter describing the choice of the prior distribution. Moreover, its performance has only been compared to the screeplot and broken-stick methods, and not to Bartlett’s test nor to the randomization procedures that performed well in previous comparative studies.

In this article, we consider several algorithms to extend and automate the Auer-Gervini method by providing objective rules to select the number of significant principal components. Using an extensive set of simulations, we compare these algorithms to the broken-stick model, Bartlett’s test, and ter Braak’s randomization methods. The methods chosen for comparison were the “winners” from the two previous comparative studies.^7,9^ Our extensions to the Auer-Gervini method are implemented in the **PCDimension** R package. Because the most promising versions of the randomization algorithms appear not to be readily available, we have also implemented them in the **PCDimension** package. For convenience, the package also implements the broken-stick method. For Bartlett’s test, we rely on an existing implementation in the **nFactors** package.

This article is organized as follows. In **Methods**, we review the theoretical framework of the four types of methods (Bartlett’s Test, the Broken-Stick method, the randomization-based procedures, and the Auer-Gervini model). In the Simulation **Study** subsection of **Results**, we perform simulation studies to test the performance of the proposed algorithms. In the **Decomposing the Apoptosis Pathway** subsection, we apply the methods to a study of apoptosis in acute myeloid leukemia (AML) using reverse phase protein arrays (RPPA). Finally we conclude the paper and make several remarks in **Conclusions**. A simple example to illustrate the implementation of the four types of methods in the **PCDimension** package is also provided in the supplementary material.

## Methods

Let *X* denote an *n* × *m* data matrix, where each row represents an object, and each column represents a measured attribute. We are primarily interested in biological applications where *n* ≪ *m*. In PCA, each principal component is a linear combination of the attributes. In this section, we briefly review four methods used to estimate the number of statistically significant PCs.

### Bartlett’s Test

Bartlett proposed a statistical method to conduct a hypothesis test on the significance of the principal components based on the eigenvalues of Σ, the correlation matrix.^8^ This test is designated to check whether the remaining eigenvalues of the correlation structure are equal after removing the well-determined (highly significant) components. Let the eigenvalues of Σ be λ_1_, …, λ_*n*_ with λ_1_ ≥ ⋯ ≥ λ_n_ ≥ 0. The procedure, for various values of *k*, starting at *n* − 2 in decreasing order, is to test the null hypothesis 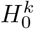 that the “(*n* − *k*) smallest eigenvalues of the correlation matrix are equal” against the alternative hypothesis 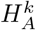 that “at least two of the (*n* − *k*) eigenvalues are different”. The test statistic is

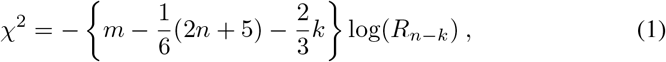

where

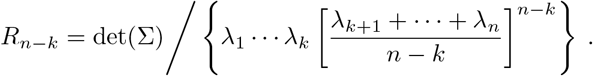

Under the null hypothesis, the test statistic follows a *χ*^2^ distribution with 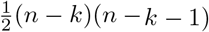 degrees of freedom. The optimal number of principal components is the smallest *k* where 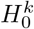 is accepted. Both Lawley^13^ and Anderson^14^ made some modifications to the multiplicative factor 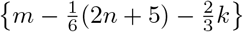 for (1); these are viewed as improved variants of Bartlett’s test.

### Broken-Stick Model

Under the assumption that the total variance of the multivariate data is divided at random among all possible components, the expected distribution of the eigenvalues in the covariance or correlation matrix follows a broken-stick distribution.^15^ This model says that, if we have a stick of unit length, broken at random into n segments, then the expected length of the *k*th longest piece is

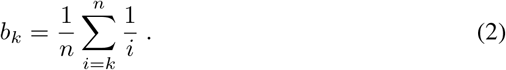

Since the expected values under the broken stick model are obtained in decreasing order, it is necessary to rank the relative proportions of the variance that are accounted for by the PCs in the same way. The estimated number of PCs is the maximum index where the observed relative proportion of variance is greater than or equal to the expected value from the broken-stick distribution.

### Randomization-Based Procedure

This procedure involves a randomization approach to generate a large number of data sets by scrambling the observed data in a manner of sampling without replacement.^10,11^ These randomized data sets are then used to compute empirical *p*-values for the statistics of interest that characterize the internal structure of the eigenvalues in the correlation matrix. The test procedure is: (1) randomize the values with all the attribute columns of the data matrix; (2) perform PCA on the scrambled data matrix; and (3) compute the test statistics. All three steps are repeated a total of (*B* − 1) times, where *B* is a large enough integer to guarantee the accuracy of estimating the *p*-value; in practice, *B* is usually set to equal 1000. In each randomization, two test statistics are computed: (1) the eigenvalue λ_*k*_ for the *k*-th principal component; and (2) a pseudo F-ratio computed as 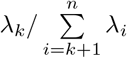. Finally, the *p*-value for each *k* and each statistic of interest is estimated to be the proportion of the test statistics in all data sets that are greater than or equal to the one in the observed data matrix.

### Auer-Gervini Model

We briefly review the Auer-Gervini method.^12^ Suppose *x_i_* ∈ ℝ^*n*^ (*i* = 1, …, *m*) are all columns of data matrix *X*, and {*x*_1_, …, *x_m_*} is a random sample with mean vector ***μ*** and covariance matrix Σ. Write Σ = ΓΛΓ^*T*^ where Γ = (λ_1_, …, γ_*n*_) is orthonormal and Λ = diag(λ_1_,…, λ_*n*_) with λ_1_ ≥ ⋯ ≥ λ_*n*_. Let 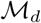 be the model with *d* significant components or eigenvalues, that is, λ_1_ ≥ ⋯ > λ_*d*_, λ_*d*_ > λ_*d*+1_, and λ_*d*+1_ = ⋯ = λ_*n*_, for *d* ≤ *n* − 1. Under 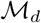, a random sample is

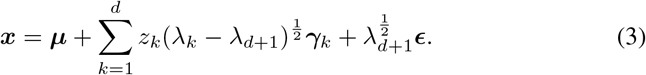

where *z*_1_, …, *z_d_* are uncorrelated random variables with mean 0 and variance 1, and *ε* is a random error vector with mean 0 and covariance matrix ***I*_n_**. Therefore the problem of selecting the number of PCs is transformed into the problem of choosing the correct model 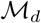.

A prior probability is assigned to 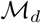 of the form

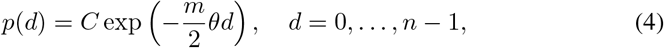

where *C* is a normalizing constant that satisfies 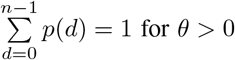. Then, under certain approximations, one can use Bayes rule to derive a formula for the maximum posterior estimate of *d* as a function of the prior parameter *θ*:

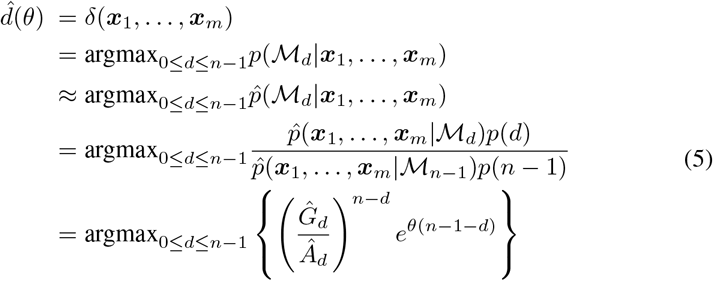

where *Ĝ_d_* and *Â_d_* are the geometric and the arithmetic means of 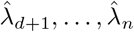, respectively. The formula describes 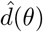 as a nonincreasing step function with respect to *θ*, with 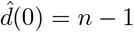. The step function is then plotted and the “highest dimension for which the step length is significantly large” is selected to be the optimal number of components.

In other words, an exponential prior is placed on the number of significant components. The prior depends on a hyperparameter *θ* ≥ 0, that governs how fast the distribution decays. As *θ* goes to ∞, the prior drops off rapidly and the posterior estimate of the number of PCs will go to 0. Auer and Gervini proposed graphing the posterior estimate as a step function of *θ*, which can visually help select the highest “nontrivial step length”. A large step length means that the estimated number of PCs is optimal under a wide range of prior model probabilities. However, the notion of “nontrivial step length” remains subjective, which is similar to the situation where one needs to select a recognizable “elbow” in the screeplot. Automating the definition of nontrivial step length is further complicated by the fact that the final step for *d* = 0 is theoretically infinite. We will operationalize the final subjective step by putting an upper bound on the largest “reasonable” estimate of *θ*, and will develop criteria to automatically choose the significantly large step length.

### Automating the Auer-Gervini Method

As originally described, the Auer-Gervini model is a visual Bayesian approach, and the critical final step is to decide what constitutes a significantly large length of a step. This problem can be thought of as one of classification, in which the set of step lengths must be partitioned into two groups (short and long). We propose to test multiple algorithms to solve the problem as follows.

#### TwiceMean

Use twice the mean of the set of step lengths as a cutoff to separate the long and short steps. Intuitively, the idea is to view the distribution of the step lengths as exponential when the data arises from random noise. Since the mean equals the standard deviation for an exponential distribution, twice the mean is the same as the mean plus one standard deviation, and provides a plausible cutoff to select “long” step lengths. This simple idea is inspired in part by Chaterjee,^16^ who considered recovery of low-rank matrices by thresholding singular values. He proposed that one could have a single universal choice of the threshold parameter which is slightly greater than 2 and gives near-optimal mean square error for Singular Value Hard Thresholding (SVHT).

#### Kmeans

Since the goal is to partition the step lengths into two groups, a natural solution is to cluster them using the traditional K-means algorithm with *K* = 2. We seed the algorithm with starting centers using the minimum and maximum step lengths. As we will discuss in more detail below, the final step (when *d* = 0) is theoretically infinite. We will bound this last step, but it will ensure that at least one step is “long”.

#### Kmeans3

Our initial experience (data not shown) using K-means with large n found it to be overly conservative when assigning steps to the “long” group. Given its dependence on Euclidean distances, that is exactly how one should expect it to perform when the true mixture distribution is skewed right. To address this problem, we modified the algorithm as follows. If the number of objects is large (*n* ≥ 55), we use the K-means algorithm, with *K* = 3 and seeds of the minimum, median, and maximum values, to separate the step lengths into three groups: Low, Intermediate, and High. We then treat both Intermediate and High groups as long.

#### Spectral

Use spectral clustering to divide the step lengths into two groups. Spectral clustering is one of the most popular modern clustering algorithms. It sometimes outperforms the traditional clustering algorithms such as K-means.^17,18^

#### CPT

Instead of simply clustering the step lengths into two groups, we can instead sort them into increasing order and view the problem as one of change point detection. One existing solution to this problem is provided by the At Most One Change (AMOC) method implemented by the `cpt.mean` function from the **changepoint** R package.^19^ The detection of the first change point is posed as a hypothesis test and a generalized likelihood ratio based approach is extended to changes in variance within normally distributed observations. The reader can consult Hinkley^20^ and Gupta and Tang^21^ for more details.

#### CpmFun

The **cpm** R package defines several other “Change Point Models”. ^22^ These are implemented by the `detectChangePointBatch` function, which processes the step lengths in one batch and returns information regarding whether the sequence contains a change point. The default is to use the “Exponential” method, which computes a generalized likelihood ratio statistic for the exponential distribution.

#### Ttest

We also implemented a novel change point detection algorithm based on the t-test. We begin by sorting the steps lengths in increasing order. Then we compute the gaps between successive step lengths. At each (sorted) step, we use the t-distribution to determine the likelihood that the next gap is larger than expected from the previously observed gaps. The first time that the next gap is significantly larger than expected, we assert that this next step length is the smallest one that constitutes a “long” step length.

#### Ttest2

Where the K-means algorithm was found to be conservative, the Ttest algorithm just described was sometimes found to be anticonservative. This can happen when the first few step lengths are all about the same size, which yields a small standard deviation. In this case, a relative short next step will be falsely discovered based on the Ttest criterion. To avoid this problem, it may be appropriate to include the next (test) step length and gap when estimating the mean and standard deviation of the gap distributon. We modified the Ttest algorithm in this way to make it more conservative.

### Bounding the Last Step

All of the proposed methods for separating short from long steps require us to bound the permissible length of the final step when 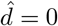, which would otherwise be infinite. This step is important, since it allows the algorithm to conclude in some cases that the only long step is the final one, and the true number of principal components should equal zero. We use the largest of the following three quantities:

1. *θ*_0_ = Three percent further than the final change point (to *d* = 0) in the step function.
2. *θ*_0_ = −2log(0.01)/*m*, where *m* is the number of attributes. This procedure selects the value of *θ* for which 99% of the prior probability is assigned to *d* = 0.
3. *θ*_0_ = (18.8402 + 1.9523 * *n* + 0.0005 * *n*^2^)/*m* if *m* ≥ n and *θ*_0_ = (18.8402 + 1.9523 * *n* + 0.0005 * *n*^2^) * *m*/*n*^2^ otherwise, where n is the number of objects. This formula was derived empirically from a Monte Carlo study on data sets with random noise. We estimate *θ*_0_ as the maximum of the empirical largest change point in the step function for various values of *m* and *n* in two scenarios: *m* ≥ *n* and *m* < *n*. It can be seen as the value where the maximum point for *d* = 0 could be achieved when various *d*’s are sharing the prior information on *θ* under the uninformative noise structure.

Note that all simulations and computations in this article were performed using version 3.2.2 of the R statistical software environment with version 1.1.3 of the **PCDimension** package, which we have developed, and version 2.3.3 of the **nFactors** package. The details on how to select the number of PCs for a simple example by using the **PCDimension** package are provided in the supplementary material.

## Results

### Simulation Study

We follow a Monte Carlo procedure to study the robustness of the four types of methods described above for estimating the number of PCs. For real data sets we will never know the “correct” answer. So, we simulate a collection of data sets with different correlation structures to compare the numerical Auer Gervini-model we have implemented with the other three types: Broken-Stick, Bartlett’s test, and the randomization-based procedure. Details about the correlation structures and data sets are provided in the next section.

#### Simulated Datatypes

We use a protocol similar to those discussed in recent papers.^9,12,23–26^ In the simulations, the number of measured attributes is taken to be either m = 100 or m = 400. The range of 100 to 400 is chosen to represent small to moderately large data sets. We also consider data sets with either *n* = 24 or *n* = 96 objects. We view 24 objects as a small data set, and 96 objects as a moderately large one. ^27^ The number of significant diagonal blocks is either the number shown in Figure 1 or twice that number (with finer correlation structures of double group blocks). By varying both the number of objects and the number of correlated blocks, we can explore the effects of the number of nontrivial components and the number of objects per component. To also explore the effects of different combinations of additional factors, including eigenvalue distributions, signed or unsigned signals, uncorrelated variation, and unskewed normal or skewed distributions, we use the 19 different covariance structures and correlation matrices illustrated in Figure 1. Matrices 1–3 are covariance matrices Σ with different marginal distributions: normal, marginal gamma, and marginal t distribution, where Σ = ΓΛΓ^*T*^, and Λ = diag(λ_1_, …, λ_*n*_) with λ_*i*_ = 1/*i* for 1 ≤ *i* ≤ 5 and λ_*i*_ = 1/10 for *i* ≥ 6. Matrices 4–19 are correlation matrices corr(*X*) with corresponding covariance matrices *σ*^2^ * corr(*X*) where *σ*^2^ = 1. Matrix 4 contains only noise; it is a purely uncorrelated structure. Matrices 5–6 represent correlation structures with various homogeneous cross-correlation strengths (unsigned signals) 0.3 and 0.8. Matrices 7–13 are correlation matrices where between-group (0.3, 0.1, or 0) and within-group (0.8, or 0.3) correlations of objects are fixed.^9,24^ Matrices 14–19 are correlation structures where negative cross-correlations (−0.8 or −0.3, signed signals) are considered within groups and mixture of signed and unsigned signals are also included.

**Figure 1.**
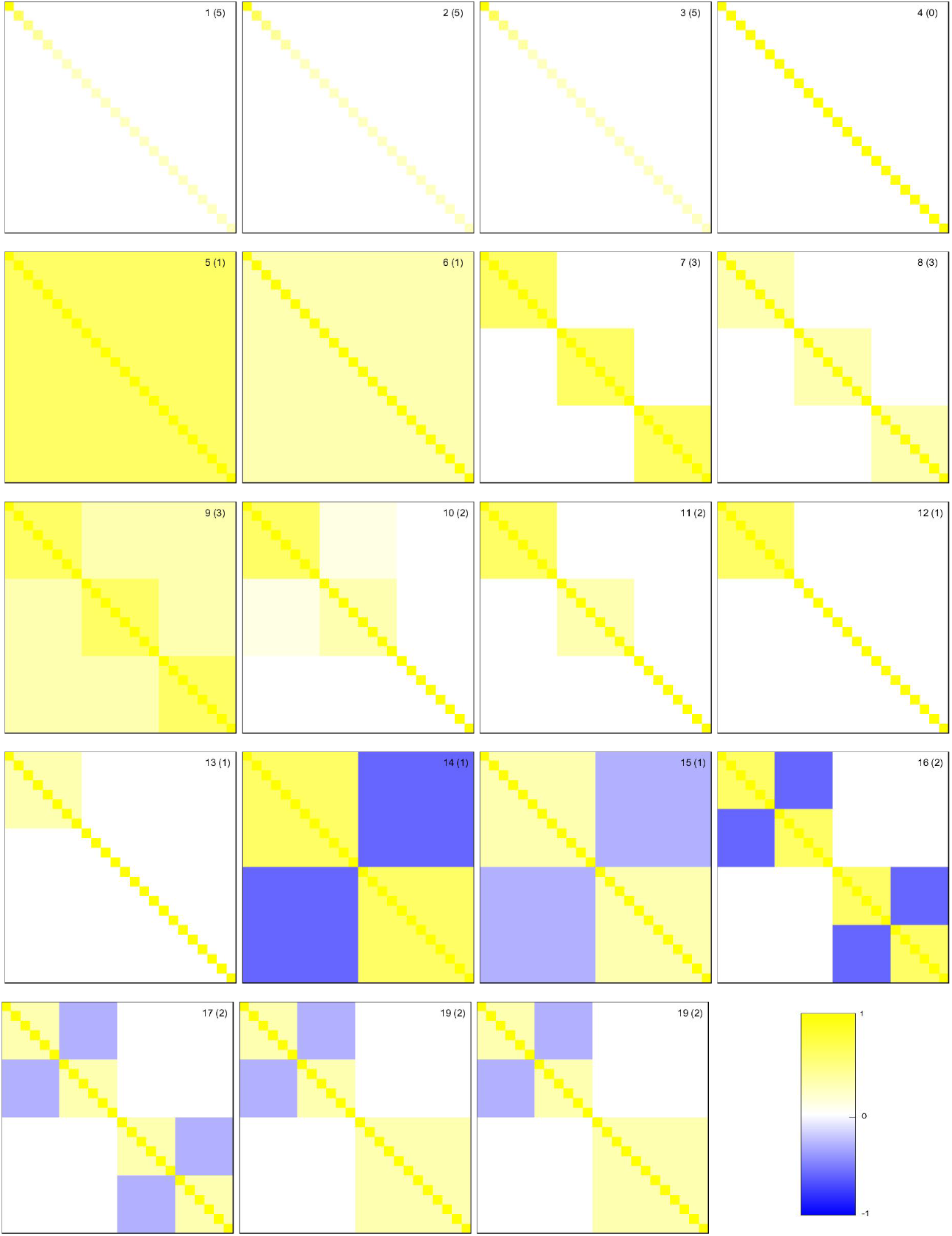
The 19 correlation matrices considered in the simulation study for 24 objects. Values of correlations are provides by the colorbar. Numbers in parentheses correspond to the known dimension.

#### Empirical Simulation Results

For each of the 19 scenarios, we simulate 1000 data sets. Then the numbers of components are estimated based on all (thirteen variants of the) four types of methods. That is, for each variant within each type of model, we compute the estimated dimension. We also investigate a “majority rule” procedure for the Auer-Gervini model; that is, the dimension that more than three criteria out of eight select is the one that is estimated by the majority rule. Then we calculate the absolute difference beween the known dimension and the estimated ones for each simulated sample and correlation structure. The mean of the absolute differences over both 1000 simulated data sets and 19 correlation matrices are plotted in Figure 2. The corresponding numeric values are also provided in Supplementary Table 1 in the Appendix. The values in this table can help assess the quality of each method. The closer to zero the values are, the better the corresponding variant within the type of method. However, the values do not describe whether a method over- or underestimates the number of non-trivial PCs.

**Figure 2.**
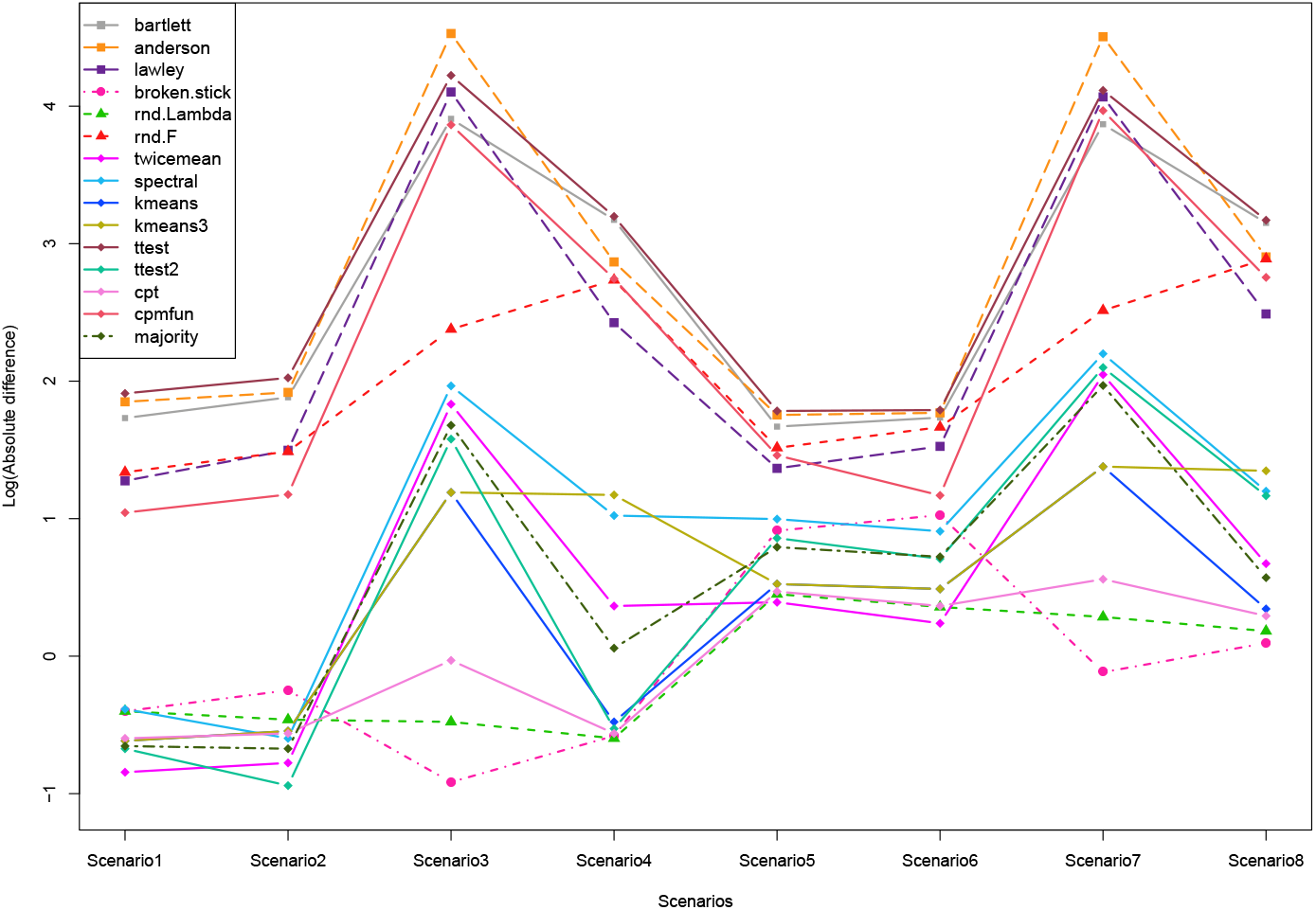
Log transformed mean values, across the correlation matrices, of the absolute difference between the known dimension and the sample estimates from 15 different methods in eight simulation scenarios (Scenario 1: 24× 100, 1X correlated blocks; Scenario 2: 24×400, 1X; Scenario 3: 96×100, 1X; Scenario 4: 96×400, 1X; Scenario 5: 24×100, 2X; Scenario 6: 24×400, 2X; Scenario 7: 96×100, 2X; Scenario 8: 96×400, 2X).

In Figure 2 and Supplementary Table 1, one can see that, as anticipated, the results from most algorithms are better with fewer correlated blocks (Scenarios 1–4), probably because there are more objects representing each block. Also, accuracy in almost all methods is better with fewer objects (Scenarios 1–2 and 5–6). The situation is more complicated when the number of attributes changes. In general, the worst performance occurs when the data matrix is nearly square (Scenarios 3 and 7).

Overall, the most accurate methods are (1) the rnd-Lambda algorithm in the randomization-based method, and (2) the Auer-Gervini model with criterion “CPT”, regardless of the matrix size or the number of blocks. They produce the most accurate results when averaged across all scenarios. However, the rnd-Lambda algorithm, whose average absolute difference is always less than one component with the original (1X) blocks and less than two with 2X blocks, only produced the best overall performance in 1 out of 8 scenarios. The results for the Auer-Gervini model with “CPT” criterion are similarly accurate, but they become worse when the number of objects (96) is close to the number of attributes (100).

By contrast, the Broken-Stick method and the “TwiceMean” criterion in the Auer-Gervini model each have the best performance for 3 out of 8 scenarios. The accuracy of the Broken-Stick method is worse when there are fewer objects, and the “TwiceMean” algorithm in the Auer-Gervini model is worse when there are more objects. These two approaches can be used as complementary ones for each other when considering the number of objects.

The variants of Bartlett’s test, as implemented in R, have the worst performance of all the stopping rules we have considered (Figure 2). Furthermore, the rnd-F algorithm is not as good as expected, since a previous study found it to be one of the most successful rules tested.^9^ Even though the “CPT” criterion is the best overall, and the “TwiceMean” criterion produces the best results for 3 out of 8 scenarios, some of the criteria that we use to automate the Auer-Gervini model are not good candidates for computing the dimension. For example, the “Ttest” and “CPM” criteria often result in large deviations of the estimated dimension as shown in Supplementary Figure 3. The “CPM” criterion does a good job in calculating the number of non-trivial components for correlation matrices 1–3, but it tends to overestimate the dimension for weakly correlated or uncorrelated matrices (matrices 4, 8, 10, 11, 12 and 13).

#### Running Time

In addition to the absolute differences between the true dimension and the estimates, we computed the average running time of the four types of methods over all correlation matrices per data set (Table 1). All timings were carried out on a computer with an “Intel® Core™ i7-4770S CPU @ 3.10 GHz” processor running Windows^®^ 8.1. Note that the time shown in the table is the total time of all the variants or criteria within each type of model. It is obvious that the computation time becomes longer as the number of objects or attributes increases, and there is almost no change in time usage when the number of blocks is doubled. From the table, we can see that Bartlett’s test and the Broken-Stick method use the least time in computing the number of components. However, the accuracy using the Broken-Stick approach is better than in Bartlett’s test as shown in Figure 2. In terms of accuracy, the “CPT” criterion in the Auer-Gervini model is comparable to the rnd-Lambda algorithm in the randomization-based method. However, the randomization-based method (using default number of 1000 permutations) spends several orders of magnitude more time than the Auer-Gervini model. To sum up, the implementation time for three type of methods, Bartlett’s test, the Broken-Stick method and the Auer-Gervini model, is negligible compared to that of the randomization-based method. At the same time, the “CPT” criterion in the Auer-Gervini method keeps the same level of accuracy as the randomization-based procedure, while Broken-Stick method is doing moderately well to some extent.

**Table 1.** Average running time of the four types of methods across correlation matrices (unit: seconds).

| Rules | Original Blocks (1X) |  |  |  | Twice Blocks (2X) |  |  |  |
| --- | --- | --- | --- | --- | --- | --- | --- | --- |
|  | 24 objects |  | 96 objects |  | 24 objects |  | 96 objects |  |
|  | m=100 | m=400 | m=100 | m=400 | m=100 | m=400 | m=100 | m=400 |
| Bartlett Test | 0.015 | 0.01 | 0.019 | 0.027 | 0.014 | 0.009 | 0.024 | 0.027 |
| Broken-Stick | 0.007 | 0.008 | 0.012 | 0.031 | 0.007 | 0.008 | 0.014 | 0.031 |
| Rand.-based | 6.858 | 8.82 | 14.562 | 35.579 | 6.906 | 8.794 | 16.622 | 35.502 |
| Auer-Gervini | 0.115 | 0.071 | 0.351 | 0.454 | 0.119 | 0.072 | 0.422 | 0.456 |

#### High-accuracy methods: rnd-Lambda, and Auer-Gervini with CPT

More detailed results on the performance of the rnd-Lambda algorithm are presented in Figure 3 and Supplementary Tables 2 and 3 in the Appendix for the case of the original block structure shown in Figure 1. Similar results for the “CPT” criterion in the Auer-Gervini model are presented in Figure 4 and Supplementary Tables 4 and 5. For each covariance or correlation matrix, we computed the percentage of deviations between the estimates and the known dimension. The results are similar regardless of the number of attributes (100 or 400) or objects (24 or 96). Both methods tend to underestimate the dimension for covariance matrices of unskewed or skewed distributions (matrices 1–3). They are quite accurate for correlation matrices of normal distribution (matrices 4–19). When they make errors with the normal distribution, the rnd-Lambda algorithm is more likely to slightly overestimate the dimension while the Auer-Gervini CPT methods is more likely to underestimate.

**Figure 3.**
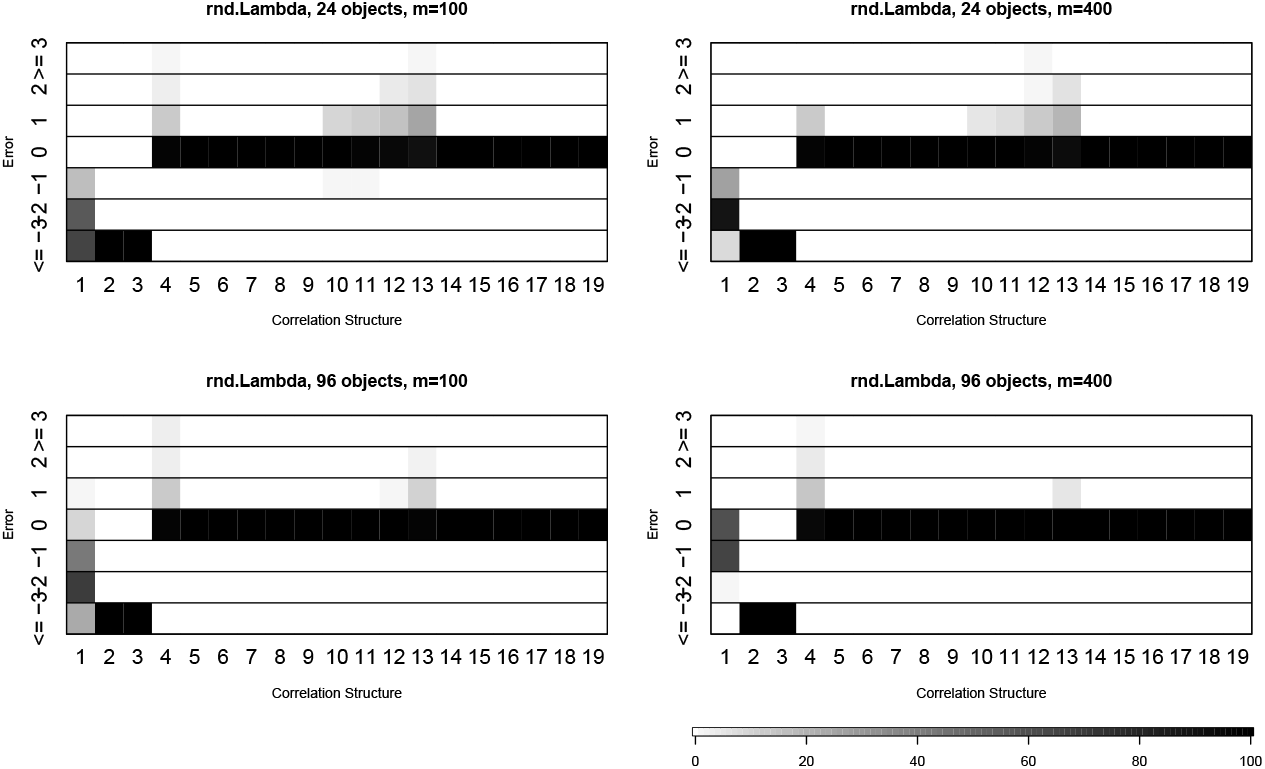
Percentage of deviations between the estimate from randomization-based procedure (rnd-Lambda) and known dimension.

**Figure 4.**
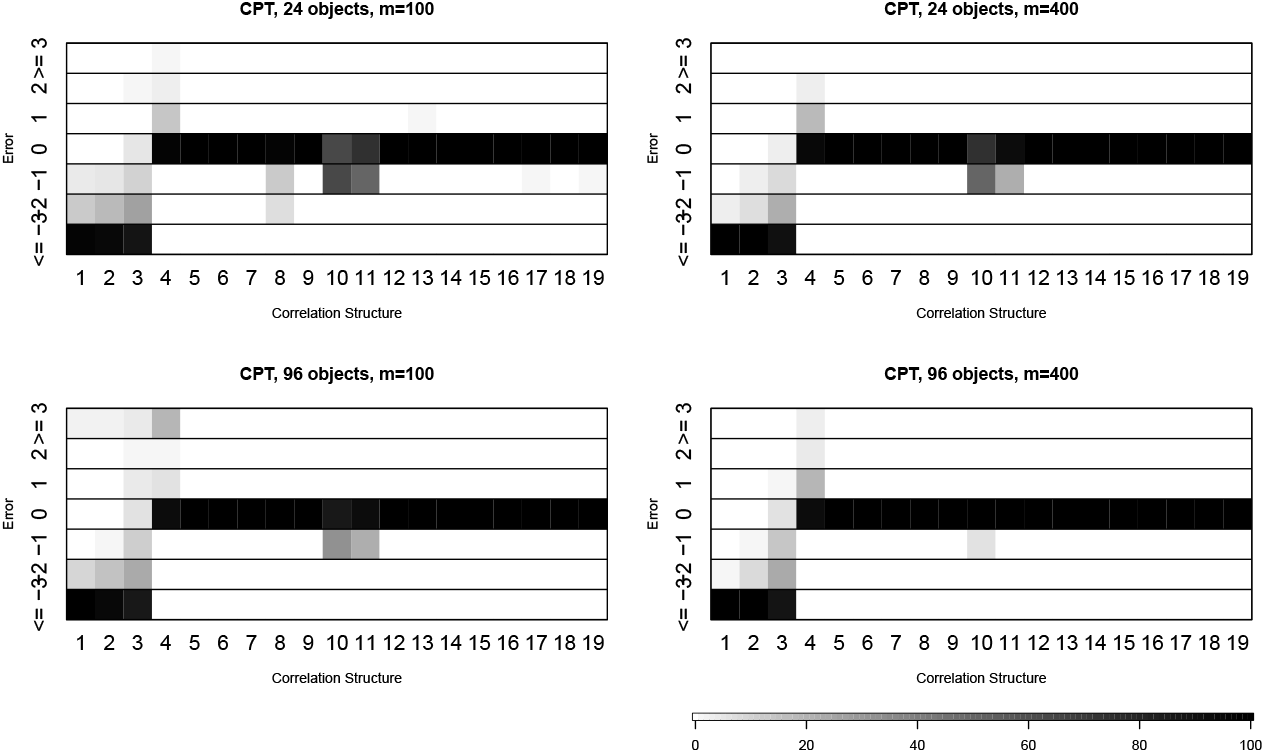
Percentage of deviations between the estimate from the “CPT” criterion in the Auer-Gervini model and known dimension.

#### High-accuracy methods: TwiceMean, and Broken-Stick

We present details on the performance of the “TwiceMean” criterion in the Auer-Gervini model and the Broken-Stick method in Figure 5 and Supplementary Tables 6 and 7. For data sets with 24 objects, the Auer-Gervini method with criterion “TwiceMean” is very accurate for uniform matrices (matrices 5–6), correlated matrices (matrices 7–13), and unsigned data with or without signed signals (matrices 14–19). When the sample covariance matrix is either skewed or unskewed (matrices 1–3), the dimension is usually underestimated. Also the results do not vary too much with different numbers of attibutes. For 96 objects, there is not much difference between the results of the Broken-Stick method in Table 7 and that of the criterion “TwiceMean” in Table 6. And the results of the Broken-Stick method for 100 attributes are slightly better than those for 400 attributes.

**Figure 5.**
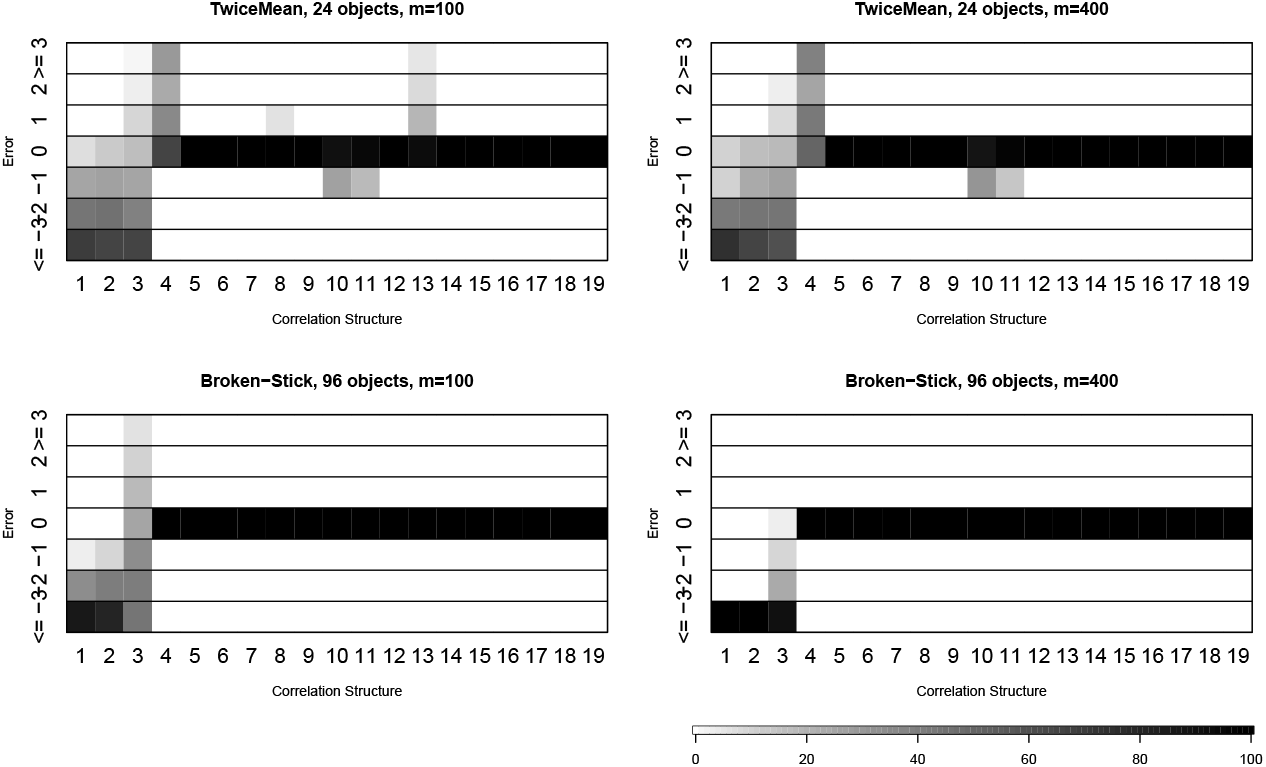
Percentage of deviations between the estimate from either “TwiceMean” criterion in the Auer-Gervin model or the Broken-Stick model and known dimension.

#### Robustness to random noise

We investigated the influence of random noise on the ability of different methods to correctly detect the underlying structure. We conducted additional simulation studies by adding different levels of (IID normal) noise corresponding to three different values of the variance: 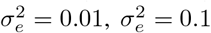 and 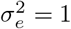. Summaries of the absolute difference between the known dimension and the esimates made using various methods under different levels of noise are presented in Supplementary Tables 8, 9, and 10. Although the error rate tends to increase with increasing noise, we found that the relative performance of the methods is consistent regardless of the size of the noise. That is, most algorithms still perform better with fewer objects, and accuracy in almost always worse when the number of blocks doubles. Most importantly, the best method for each scenario doesn’t change in most cases as the noise moves from 0 to 1. The “TwiceMean” criterion in the Auer-Gervini model is better when the number of objects is small relative to the number of objects, while the Broken-Stick method is better when the number of objects is close to the number of attributes. Finally, regardless of the noise level, the rnd-Lambda algorithm and the Auer-Gervini model with “CPT” criterion outperform the others on average across all scenarios.

### Decomposing the Apoptosis Pathway in AML

Since the introduction of gene expression microarrays in the 1990s, most statistical analyses of omics data have treated pathways as second-class objects, in the following sense: primary analyses are performed at the gene level. That is, the data is first analyzed gene-by-gene to find differences between known groups of patients such as responders and non-responders. Then a significance cutoff is chosen and a second statistical test conditional on the gene-by-gene results (like gene set enrichment analysis^28^) is performed in order to infer which pathways differ between the two groups. One reason analysts give precedence to individual genes is that univariate analyses are easier than the multivariate ones needed for pathways. However, many biologists are more interested in pathways than in individual genes, because they give a higher-level functional picture of biological behavior.

Informally, biologists talk about pathways as though they are one-dimensional entities. At a cell level they are “on” or “off”; at a tissue level, they have a simple “degree of activation”. But we hypothesize that most pathways, including the apoptosis signaling pathway, are intrinsically multidimensional. To test this hypothesis, we used a subset of reverse phase protein array (RPPA) data on samples collected from 511 patients with acute myeloid leukemia.^29,30^ The subset consists of 33 proteins that are involved in the apoptosis signaling pathway. Apoptosis is known to be an essential component of several processes including normal cell turnover, proper development and functioning of the immune system, and chemical-induced cell death.^3^ It is generally characterized by distinct morphological states and energy-dependent biochemical mechanisms. Even though many important apoptotic proteins have been identified, the molecular mechanisms of these proteins still remain to be elucidated.

We applied 14 different methods to determine the number *d* of significant components in this RPPA data set; the results are displayed in Table 2. The number of components that they find is highly variable, ranging from *d* =1 to 31. Even the four best methods from our simulations give different values (Broken-Stick: *d* =1; CPM: *d* =1; TwiceMean: *d* =6, and rnd-Lambda: *d* = 8). However, our simulation studies suggest that the TwiceMean Auer-Gervini method works particularly well when the number of objects (in this case, 33 proteins) is relatively small. In the top panel of Figure 6, we have plotted the maximum posterior estimate of the number of components as a function of the prior parameter *θ*; this plot gives better support for *d* = 6 than for *d* =8.

**Figure 6.**
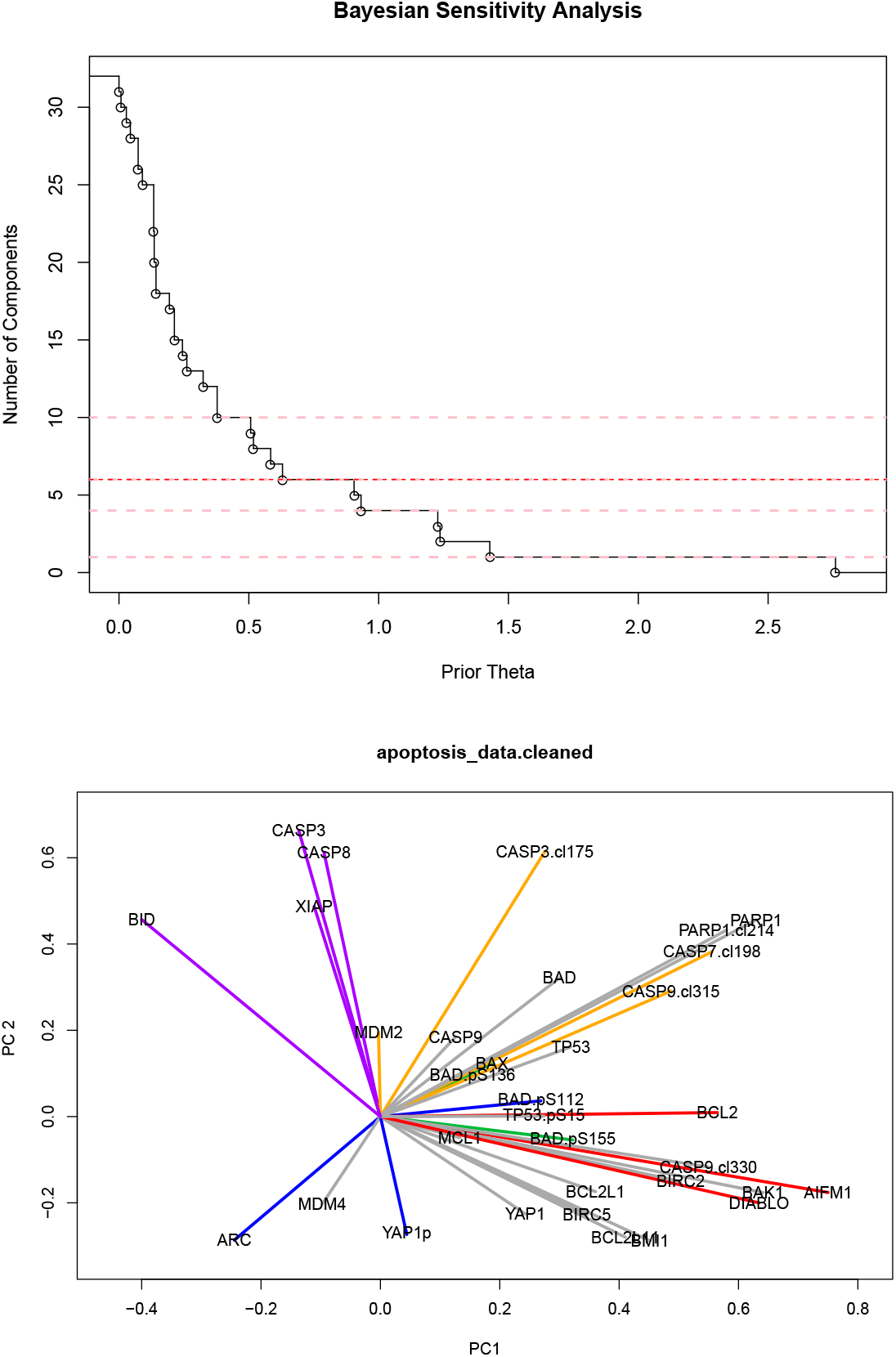
Analysis of AML RPPA data. (Top) Auer-Gervini step function relating the prior hyperparameter *θ* to the maximum posterior estimate of the number 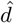 of significant principal components. (Bottom) Projection of proteins on the space of the first two components; colors denote different clusters.

**Table 2.**
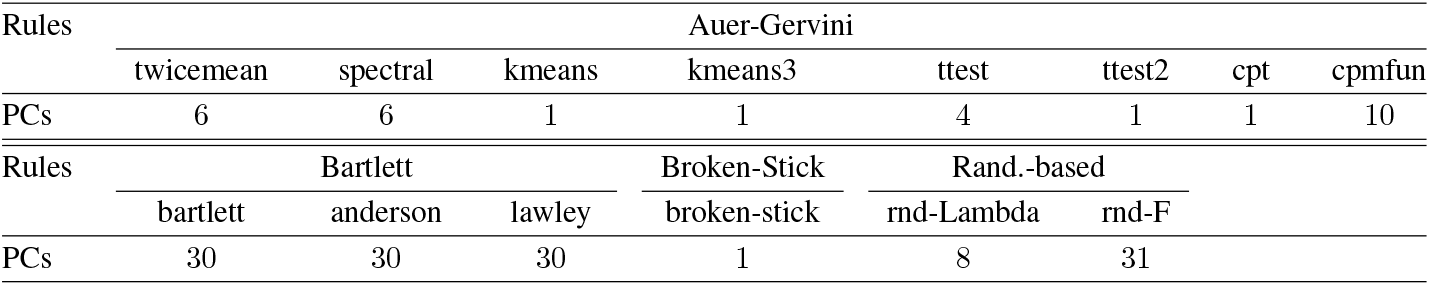
Number of principal components (PCs) from different algorithms on RPPA data

In order to understand the biology driving these mathematical principal components, we then projected the proteins into the 6-dimensional PC space and used their directions to cluster them using a von-Mises Fisher mixture model.^31^ Using the Bayesian Information Criterion for model selection, we found an optimal clustering into 6 groups of proteins, which are displayed in different colors in a 2-dimensional projection in the bottom panel of Figure 6. The six protein clusters are:

1. AIFM1, BCL2, and DIABLO, which we interpret as a block corresponding to the mitochondrial release of apoptosis-inducing factor^32^;
2. BID, CASP3, CASP8, and XIAP, which are part of a caspase-3 feedback loop^33^;
3. CASP3.cl175, CASP7.cl198, CASP9.cl315, and MDM2, a cleaved caspase block^34^;
4. ARC, BAD.pS112, and YAP1p;
5. BAD.pS136, BAD.pS155, and BAX; and
6. the core group of 16 apoptosis-related proteins: BAD, BAK1, BCL2L1, BCL2L11, BIRC2, BIRC5, BMI1, CASP9, CASP9.cl330, MCL1, MDM4, PARP1, PARP1.cl214, TP53, TP53.pS15 and YAP1.

The fact that three of these six clusters can be immediately identified from the literature as coherent biological subcomponents of the apoptosis pathway provides strong support for our approach.

## Conclusion

PCA is one of the most popular and important techniques in the multivariate analysis of general data sets. However, because principal components are linear combinations of correlated variables, these components usually lack interpretability when analyzing biological data sets, especially transcriptomic or proteomic data sets from cancer patients. Our study of the apoptosis pathway using proteomic data from AML patients shows that we can use mixture model clustering in principal component space to replace the uninterpretable mathematical components with natural collections of related genes that enhance the biological interpetability of the decomposition. We expect that applying these methods to the genes or proteins in other signaling pathways will divide these pathways into one-dimensional “building blocks” that are interpretable, robust, and can yield new biological insights. It would be of particular interest to apply these ideas to overlapping pathways to better understand the way similar components are reused in different contexts.

Our ability to find biologically interpretable components, however, depends in a fundamental way on being able to determine the dimension of the principal component space. To accomplish this task, we introduced the **PCDimension** R package, which implements three types of models—the Broken-Stick method, the randomization-based procedure of ter Braak,^10,11^ and our enhancments to the model developed by Auer and Gervini^12^—to compute the number of significant principal components. Through extensive simulations, we have shown that the enhanced Auer-Gervini methods are competitive with the methods that performed best in previous comparative studies.

It has been claimed that simulation of multivariate data sets can always be criticized as unrepresentative, since they can never explore more than a tiny fraction of the wide range of possible covariance and correlation structures^4^ As with previous simulation studies, our work may suffer from the same limitation. However, Ferre has also pointed out that simulations are the only way to test and compare these methods.^7^ It is still valuable to compare methods empirically when the dimension of the data set is known, and factors of interest can be manipulated under simulation. We have endeavored to explore a wide variety of different correlation structures in order to identify settings where each method is likely to fail.

In our simulations, the variants of Bartlett method clearly had the worst performance. This finding may be somewhat surprising: Peres-Neto and colleagues^9^ found these methods to be only a little worse than the best performers in their simulations and concluded that they were actually the best for distinguishing *d* = 0 from *d* ≥ 1. There are two factors that distinguish their simulations from ours. First, we considered a wider variety of correlation structures. The matrices considered by Peres-Neto are represented by our matrices 4–13. Second, the matrices we consider are larger. Motivated by problems from ecology, they looked at matrices that were 9 × (30 or 50) or 18 × (60 or 100). Motivated by the larger gene or protein expression data sets currently being produced in biology, we looked at matrices that were (24 or 96) × (100 or 400). We think that both factors contribute to the different results. Supplementary Figure 3 in the Appendix shows that the errors arise primarily from matrices 1–4 and 10–12. This subset is comprised precisely of those matrices that include a substantial amount of unstructured noise. We suspect that the underlying difficulty arises because the stepwise hypothesis tests give rise to a classical problem of multiple comparisons. Thus the likelihood of incorrectly rejecting a null hypothesis (and inflating the dimension) increases. This may also explain why Ferre^7^ cautioned that Bartlett’s test can overestimate the number of components.

The rnd-F method introduced by ter Brack also performed significantly worse in our simulations than in those of Perres-Neto. The plots in Supplementary Figure 3 reveal two things. First, for virtually every correlation matrix we considered, rnd-F is more variable and less accurate than rnd-Lambda. Second, the most serious large errors arise from matrices 10–13. These matrices contain a mix of both highly structured data and completely unstructured noise. Similar matrices were considered in the previous study, so we conclude that the rnd-F method simply works poorly with larger matrices.

We investigated the graphical Bayesian method of Auer-Gervini^12^ in some detail. Specifially, we introduced and tested eight algorithms to enhance the method by automatically selecting the number of components. Two of these—the novel T-test-based changepoint algorithm and the exponential model from the **CPM** package—were abysmal. In virtually every simulation, the T-test seriously overestimates the number of components. The exponential CPM model overestimates the number at least half the time. The remaining six methods have at least acceptable performance most of the time.

The clear overall winners from our simulation study are the rnd-Lambda method among the randomization-based procedures and the Auer-Gervini model with criterion “CPT”. They produce the best results on average across all 19 correlation matrices and data sets of different sizes. It is interesting to note, however, that they rarely give the absolute best results for any fixed size of data matrix. Their ultimate strength is their consistency: their estimates are always competitive with the best methods, and the average error in the estimated dimension is always less than two.

There is little to choose between the two winning methods in terms of accuracy. However, there is a clear difference in computation time. The “CPT” criterion in the Auer-Gervini model is at least two orders of magnitude faster than the rnd-Lambda algorithm, with no loss in accuracy. Not surprisingly, computation time increases for all methods as the number of objects or attributes increases. Changing the number of correlated blocks has little effect on computation time. The fastest methods overall are the variants of Bartlett’s test and the Broken-Stick method, but this increase in speed is obtained at a much bigger cost in accuracy.

Two other methods perform well in complementary settings. Both the Broken-Stick model and the Auer-Gervini method using the TwiceMean criterion are the most accurate on average for three of the eight scenarios we considered. All three cases where TwiceMean wins have 24 objects; all three cases where Broken-Stick wins have 96 objects. The Twice Mean criterion appears to severely overestimate the number of components when the size of the data matrix increases. By contrast, the Broken-Stick model appears to benefit from having extra data available. MacArthur^15^ and De Vita^35^ showed that the Broken-Stick model worked well when fitting the relative abundance data of species in ecological populations. It is possible that the distribution of the expected lengths in Equation (2) will be better approximated with larger data matrices. This may explains why its performance improves.

Our simulation studies uncovered at least two (possibly related) contexts where it is particularly difficult to estimate the number of components correctly. Every reasonable method (that is, not Bartlett’s, rnd-F, nor Auer-Gervini with the simple T-test or the exponential CPM criteria) severely underestimates the dimension for correlation matrices 1–3. In addition, most methods overestimate the dimension when there are 96 objects and 100 attributes, especially when we doubled the number of correlated blocks. In both cases, there is very little redundancy in the signals we are trying to detect. (The biological contexts where we expect to apply these methods are expected to contain considerable redundancy in the form of highly correlated genes or proteins.) To handle more general data sets, however, new methods will need to be developed in order to improve performance in these examples without sacrificing it in other examples. Further work will also be needed to clarify what kinds of structural changes occur as the number of objects increases from 24 to 96 and beyond.

We do not expect our study to be the final word on how to determine the number of significant principal components; like Ferre^7^, we must conclude that there is no ideal solution to the problem. If forced to choose one method for all data sets, we would pick the Auer-Gervini model using the CPT criterion, since it is both reasonably accurate and reasonably fast. This would especially be the case if we were trying to analyze many data sets at once; for example, when performing an analysis like the one in our AML data set for a long list of different biological pathways or gene sets. When focused on only one data set, we would compute the estimates from other Auer-Gervini enhancements (TwiceMean, Kmeans3, and Ttest2), as well as the Broken-Stick and rnd-Lambda methods, and review them in light of both the traditional screeplot and the Auer-Gervini plot of the maximum posterior dimension as a function of the hyperparameter. We believe that this combination of analytical and graphical methods, as provided in the **PCDimension** package, will guide researchers to the most reliable results.

## Funding

This project is supported by NIH/NCI grants P30 CA016058 and R01 CA182905.

